## Supplementary Material for "A biofilm-tropic *Pseudomonas aeruginosa* bacteriophage uses the exopolysaccharide Psl as receptor"

**Table S1. Strains and Plasmids**

| Strain | genotype | Reference |
|---|---|---|
| RP1860 | PAO1F wild type | (1) |
| RP44 | PA14 wild type | (2) |
| RP4708 | PAO1F Δ*fliF* | this study |
| RP4126 | PAO1F Δ*pilA* | this study |
| RP9170 | PAO1F Δ*fliF* Δ*pilA* | this study |
| RP12980 | PAO1F Δ*fliF2* | this study |
| RP13077 | PAO1F Δ*pslC* | this study |
| RP13080 | PAO1F Δ*fliF2* Δ*pslC* | this study |
| RP13081 | PAO1F Δ*fliF2* Δ*pslD* | this study |
| RP13089 | PAO1F Δ*wspF* | this study |
| RP3302 | PAO1F Δ*fliC* | (3) |
| RP12922 | PAO1F Δ*fliE* | this study |
| RP12894 | PAO1F Δ*fliG* | this study |
| RP12935 | PAO1F Δ*flgBCDEFG* | this study |
| RP12859 | PAO1F Δ*flgI* | this study |
| RP12936 | PAO1F *flhA(R147A)* | this study |
| RP12897 | PAO1F Δ*cheA* | this study |
| RP12896 | PAO1F Δ*fliHIJ* | this study |
| RP12923 | PAO1F Δ*flhA* | this study |
| RP12911 | PAO1F Δ*fliOPQRflhB* | this study |
| RP1865 | PAO1F Δ*fleQ* | (4) |
| RP13249 | PAO1F Δ*fliF2* Δ*fleQ* | this study |
| RP12961 | PAO1F Δ*fliF* Δ*gmd* | this study |
| RP12963 | PAO1F Δ*fliF* Δ*wbpM* | this study |
| RP13195 | PAO1F Δ*fliF2* Δ*ssg* | this study |
| DH5α | F– *endA1 glnV44 thi-1 recA1 relA1 gyrA96 deoR nupG purB20 φ80dlacZΔM15 Δ(lacZYA-argF)U169, hsdR17(rK–mK+), λ–* | |
| SM10 λ*pir* | *thi thr leu tonA lacY supE recA*::RP-2Tc::Mu *kanR*::λ*pir* | (5) |

| plasmid | genotype | Reference |
|---|---|---|
| pPSV37 | *colE1* origin, *gentR*, PA origin, *oriT*, $p_{lacUV5}$ promoter, *lacIq* | (6) |
| pP37-*fliF* | *fliF* under control of $p_{lacUV5}$ promoter | this study |
| pJN105 | *gentR*, $p_{BAD}$ promoter, *araC*, *pBBR1* origin | (7) |
| pJN2133 | pJN105 with PA2133 under control of $p_{BAD}$ | (8) |
| pP25-GFPo | constitutively expressed GFPmut3, *bla* | (9) |
| pPSV35-CV | *colE1* origin, *gentR*, PA origin, *oriT*, $p_{lacUV5}$, *lacIq,VSV-G tag* | (10) |
| pPSV35-CV-*pslC* | *pslC-VSV-G* under control of $p_{lacUV5}$ promoter | Joseph Mougous, unpublished |
| pPSV35-CV-*pslD* | *pslD-VSV-G* under control of $p_{lacUV5}$ promoter | Joseph Mougous, unpublished |
| pEXG2 | allelic exchange vector, *colE1* origin, *oriT*, *gentR*, *sacB* | (11) |
| pEXG2-Δ*pilA* | *pilA* deletion removing codons 3-148, in pEXG2 | this study |
| pEXG2-Δ*fliF* | *fliF* deletion removing codons 3-597, in pEXG2 | this study |
| pEXG2-Δ*fliF2* | *fliF* deletion removing codons 65-490, in pEXG2 | this study |
| pEXG2-Δ*pslC* | *pslC* deletion removing codons 8-297, in pEXG2 | this study |
| pEXG2-Δ*pslD* | *pslD* deletion removing codons 14-247, in pEXG2 | this study |
| pEXG2-Δ*fliE* | *fliE* deletion removing codons 3-108, in pEXG2 | this study |
| pEXG2-Δ*fliG* | *fliG* deletion removing codons 1-339, in pEXG2 | this study |
| pEXG2-Δ*fliHIJ* | *fliHIJ,* clean deletion moving *fliJ* STOP to *fliG*, in pEXG2 | this study |
| pEXG2-Δ*fliOPQRflhB* | deletes *fliO*(codon 2)-*flhB*(codon 377), in pEXG2 | this study |
| pEXG2-Δ*flgBCDEFG* | deletes *flgB*(codon 1)-*flgG*(codon 253), in pEXG2 | this study |
| pEXG2-Δ*flgI* | *flgI* deletion removing codons 3-368, in pEXG2 | this study |
| pEXG2-Δ*flhA* | *flhA* deletion removing codons 4-704, in pEXG2 | this study |
| pEXG2-*flhA(R147A)* | *flhA* R147A mutation, in pEXG2 | this study |
| pEXG2-Δ*fleQ* | *fleQ* deletion removing codons 4–487, in pEXG2 | (4) |
| pEXG2-Δ*wspF* | *wspF* deletion removing codons 11-329, in pEXG2 | this study |
| pEXG2-Δ*cheA* | *cheA* deletion removing codons 3-752, in pEXG2 | this study |

| | | |
|---|---|---|
| pEXG2-Δ*gmd* | Δ*gmd*, clean deletion moving *rmd* stop to *gmd* stop and mutating ATG of gmd to GTG | this study |
| pEXG2-Δ*wbpM* | *wbpM* deletion removing codons 3-646, in pEXG2 | this study |
| pEXG2-Δ*ssg* | *ssg* deletion, in pEXG2 | this study |

**Table S2. Primers**

| Name | Sequence (5'->3') | use |
|---|---|---|
| fliF-5-1 | AAAAAGAATTCACGGACCCTGCAGCGGTCGCGGGCAT | Δ*fliF*, EcoRI |
| fliF-5-2 | AACTCGAGCCGCAAGCATGCTGAAGGCCATGAACTAGTTATCCTCGCGCCGCT | |
| fliF-3-1 | TTCAGCATGCTTGCGGCTCGAGTTGAGTAAGCGCCATGAGTGAGAATCGT | |
| fliF-3-2 | AAAAAAAGCTTGTTCAGCGAGGAGACGCGCAGGACGAT | Δ*fliF*, HindIII |
| fliF2-5-2 | CACGCCGAGCACCTGCTTGACGATCTGCTGGGACCAGAGCACGACG | Δ*fliF2* |
| fliF2-3-1 | GTCGTGCTCTGGTCCCAGCAGATCGTCAAGCAGGTGCTCGGCGTG | |
| FliF-5G | TGACCATGATTACGAATTGTAGCTAGCTAGGAATTCCGGCGCGAGGATAACTAGTTCATG | Gibson clone *fliF* |
| FliF-3G | CCCGTTTAGAGGCCCCAAGGGGTTATGCTAAAGCTTTTACTCATCGGCGTTGATCCACTC | into pPSV37 EcoRI/HindIII |
| pilA-5-1 | TTCAGCATGCTTGCGGCTCGAGTTTTTCATGAATCTCTCCGTTGATTAT | |
| pilA-5-2 | AAAAAGAATTCGCGGGGTCGGAGATGCCTACAAA | Δ*pilA*, EcoRI |
| pilA-3-1 | AAAAATCTAGACGGTATCGACCGGGCAATTGCCGA | Δ*pilA*, XbaI |
| pilA-3-2 | AACTCGAGCCGCAAGCATGCTGAAAACTAAGGTGATCGAAGGTGGCTT | |
| pslC-5-1 | GCACGATCATGCGCACCCGTGGAAATTAATTAAGGTACCCGCGCAGCCAGGACGTCAAG | Δ*pslC*, |
| pslC-5-2 | CTTCCAGTAGCCTGGAAACAGGGAATGACCAGGGCGCAGC | Gibson clone |
| pslC-3-1 | GCTGCGCCCTGGTCATTCCCTGTTTCCAGGCTACTGGAAG | into pEXG2 |
| pslC-3-2 | TTATACGAGCCGGAAGCATAAATGTAAAGCAAGCTTGGATCGCCTGCTCCATGTTG | KpnI/HindIII |
| pslD-5-1 | ACGATCATGCGCACCCGTGGAAATTAATTAAGGTACCCCTCGGAACTGGGGATCAAGACC | Δ*pslD*, |
| pslD-5-2 | CATTGTTGACGGTGTAGCTGACCAGGGCGAGCATGGCGAGC | Gibson clone |
| pslD-3-1 | CTCGCCATGCTCGCCCTGGTCAGCTACACCGTCAACAATG | into pEXG2 |
| pslD-3-2 | TTATACGAGCCGGAAGCATAAATGTAAAGCAAGCTTGGTGAAGCTGATCTCCATCAC | KpnI/HindIII |
| fliE-5-1 | CGATCATGCGCACCCGTGGAAATTAATTAAGGTACCCACCCCGCCGAGCGTCGAGATTCC | Δ*fliE*, |
| fliE-5-2 | CTAGTTATCCTCGCGCCGCTCAGACACTCATGACTCTTCCTCCAACAGCC | Gibson clone |
| fliE-3-1 | GGCTGTTGGAGGAAGAGTCATGAGTGTCTGAGCGGCGCGAGGATAACTAG | into pEXG2 |
| fliE-3-2 | ATTATACGAGCCGGAAGCATAAATGTAAAGCAAGCTTCGAAACCGACGTTGTTGTCGGTC | KpnI/HindIII |
| fliG-5-1 | GATCATGCGCACCCGTGGAAATTAATTAAGGTACCGAAGACCGGCGAGGTCAGCCACCAG | Δ*fliG*, |
| fliG-5-2 | CTTTGTCCTTGTCGTGGGGGACCATTACTCATCGGCGTTGATCCAC | Gibson clone |
| fliG-3-1 | GTGGATCAACGCCGATGAGTAATGGTCCCCCACGACAAGGACAAAG | into pEXG2 |
| fliG-3-2 | TTATACGAGCCGGAAGCATAAATGTAAAGCAAGCTTCGCCCATCGGCAGCAGCTTCAGAG | KpnI/HindIII |
| fliH-5-1 | GATCATGCGCACCCGTGGAAATTAATTAAGGTACCGTCTCCTCGCTGAACACCGTGCAG | Δ*fliHIJ*, |
| fliH-5-2 | GTCGCGTCCGTTCTTCTGAGTCGTCAGATCATCTCCTCGCCACCCTTG | |
| fliJ-3-1 | CAAGGGTGGCGAGGAGATGATCTGACGACTCAGAAGAACGGACGCGAC | |
| fliJ-3-2 | ATACGAGCCGGAAGCATAAATGTAAAGCAAGCTTGTGGCTGGTACATCGGCAACTGGACC | |
| fliO-5-1 | CGATCATGCGCACCCGTGGAAATTAATTAAGGTACCTGGCGCTACAGATTCTCGAAGC | Δ*fliOPQRflhB*, |
| fliO-5-2 | GCGTAGCTCCAGCCAGGTCACTCCGCATGTCAGCGCAGCTTCTTGATG | Gibson clone |
| flhB-3-1 | CATCAAGAAGCTGCGCTGACATGCGGAGTGACCTGGCTGGAGCTACGC | into pEXG2 |
| flhB-3-2 | ATTATACGAGCCGGAAGCATAAATGTAAAGCAAGCTTGATCCGTCCGGCGACCACATCGG | KpnI/HindIII |
| flgB-5-1 | AGCACGATCATGCGCACCCGTGGAAATTAATTAAGGTACCTCCGGCGCCTATGTGCACTTCGACATG | Δ*flgBCDEFG,* |
| flgB-5-2 | CAAAGATTCTGGGTGACGAAGGACAAGGCTGCGAACCTGTTTGCCGGTGTG | Gibson clone |

| | | |
|---|---|---|
| flgG-3-1 | CACACCGGCAAACAGGTTCGCAGCCTTGTCCTTCGTCACCCAGAATCTTTG | into pEXG2 |
| flgG-3-2 | CACATTATACGAGCCGGAAGCATAAATGTAAAGCAAGCTTGATCGAACCGGTCAGGCTGTTGCTCTGC | KpnI/HindIII |
| flgI-5-1 | AGCACGATCATGCGCACCCGTGGAAATTAATTAAGGTACCCTTCCGCGTGGGCGACATCATCAC | Δ*flgI*, |
| flgI-5-2 | GAATCCATGGCGTCGTCCTCAAATGGTCATCGCGAGCGTCCTCAGAAC | Gibson clone |
| flgI-3-1 | GTTCTGAGGACGCTCGCGATGACCATTTGAGGACGACGCCATGGATTC | into pEXG2 |
| flgI-3-2 | CACATTATACGAGCCGGAAGCATAAATGTAAAGCAAGCTTAGGCGCCTCTGGTTGAGCAGCTTC | KpnI/HindIII |
| flhA-5-1 | CGATCATGCGCACCCGTGGAAATTAATTAAGGTACCCCGCTTTCGCTGGGACTGAGCCTG | Δ*flhA*, |
| flhA-5-2 | CGCAGCCCTCGTTCAGTTCTGTCCGCGATCCACTCTCGACTCCCCTGC | Gibson clone |
| flhA-3-1 | GCAGGGGAGTCGAGAGTGGATCGCGGACAGAACTGAACGAGGGCTGCG | into pEXG2 |
| flhA-3-2 | TATACGAGCCGGAAGCATAAATGTAAAGCAAGCTTGGCCCCAGGCGATGGAACCCAGTTG | KpnI/HindIII |
| flhAR147A-5-1 | GATCATGCGCACCCGTGGAAATTAATTAAGGTACCCGCGCATTCCGGCGTCAAAAG | *flhA(R147A)*, |
| flhAR147A-5-2 | GAAGCGCGCGCTGACTTCGGAAATGGCCCCGGCGCCCTTGGTCACCACCAC | Gibson clone |
| flhAR147A-3-1 | GTGGTGACCAAGGGCGCCGGGgccATTTCCGAAGTCAGCGCGCGCTTCAC | into pEXG2 |
| flhAR147A-3-2 | TTATACGAGCCGGAAGCATAAATGTAAAGCAAGCTTGCCGAGGCCGATGAAGGAAACATG | KpnI/HindIII |
| flhAR147A-Test | CGCGCTGACTTCGGAAATggc | + with 5-1 primer and mutation |
| wspF-5-1 | GCACGATCATGCGCACCCGTGGAAATTAATTAAGGTACCCCGGCGCCTTGCTGGACGACG | Δ*wspF*, |
| wspF-5-2 | CTAATCGAATACCTCCGCCAGCGGCATGTCATTGACGATTCC | Gibson clone |
| wspF-3-1 | GGAATCGTCAATGACATGCCGCTGGCGGAGGTATTCGATTAG | into pEXG2 |
| wspF-3-2 | TTATACGAGCCGGAAGCATAAATGTAAAGCAAGCTTGATGGCGTCCGGCAGCTTGACC | KpnI/HindIII |
| cheA-5-1 | GATCATGCGCACCCGTGGAAATTAATTAAGGTACCCAGTTGTCCTCGCAGCTCAATGAC | Δ*cheA*, |
| cheA-5-2 | CGGAAACCCATACGCGGCGTCAGATGCTCATTCGGCTGCTCCCAGAGACGTGTTAC | Gibson clone |
| cheA-3-1 | GTAACACGTCTCTGGGAGCAGCCGAATGAGCATCTGACGCCGCGTATGGGTTTCCG | into pEXG2 |
| cheA-3-2 | ATACGAGCCGGAAGCATAAATGTAAAGCAAGCTTCATGGCTGGACGAGGTGGCCGAAG | KpnI/HindIII |
| gmd-3-1 | ATTATACGAGCCGGAAGCATAAATGTAAAGCAAGCTTCTGGTCGCCCAGCTCGATATAGC | Δ*gmd* |
| gmd-3-2 | GACTGGGAGTCACGGGTACGAGAAGAGTGAGCCATGCTGATTCCCGTGGTGCTTTCCGGC | Gibson clone |
| gmd-5-1 | GCCGGAAAGCACCACGGGAATCAGCATGGCTCACTCTTCTCGTACCCGTGACTCCCAGTC | into pEXG2 |
| gmd-5-2 | GACAGGAGCACGATCATGCGCACCCGTGGAAATTAATTAAGAGTCGCTGTGCCTGCAGTG | no A-band LPS (12) |
| wbpM-3-1 | ATTATACGAGCCGGAAGCATAAATGTAAAGCAAGCTTCTGCGCGCCCGGTTCTTCTCCAG | Δ*wbpM* |
| wbpM-3-2 | CTATTGAACGGGGCTGATAAATAGGATGTTGTATGCGCCTGACGGTGAAATCGTCGACTG | Gibson clone |
| wbpM-5-1 | CAGTCGACGATTTCACCGTCAGGCGCATACAACATCCTATTTATCAGCCCCGTTCAATAG | into pEXG2 |
| wbpM-5-2 | GACAGGAGCACGATCATGCGCACCCGTGGAAATTAATTAATAGGTATCGACGGTGCTGTG | No B-band LPS (13) |
| PA5001-5-1 | GCACGATCATGCGCACCCGTGGAAATTAATTAAGGTACCCATTTCGATCCGCACCCGGAC | Δ*ssg* |
| PA5001-5-2 | CGTCGCCAGGTCTTCTCCAGCTCTTTCTGGACCAGAAACAGAAC | Gibson clone |
| PA5001-3-1 | CTGTTTCTGGTCCAGAAAGAGCTGGAGAAGACCTGGCGACG | into pEXG2 |
| PA5001-3-2 | TTATACGAGCCGGAAGCATAAATGTAAAGCAAGCTTCACCTGTTTCTCCAGGCGTTGC | LPS outer core mutant (14) |
| Mar1x | GGGAATCATTTGAAGGTTGGTAC | 1st rnd. TnSeq |
| olj376 | GTGACTGGAGTTCAGACGTGTGCTCTTCCGATCTGGGGGGGGGGGGGGGG | 1st rnd. TnSeq |
| Mar2-InSeq | AATGATACGGCGACCACCGAGATCTACACCATTTAATACTAGCGACGCCATCTATGTGTCAG | 2nd rnd. TnSeq |

| | | |
|---|---|---|
| TdT_Index_1 | CAAGCAGAAGACGGCATACGAGATCGTGATGTGACTGGAGTTCAGACGTGTGC TCTTCCGATCT | 2nd rnd. TnSeq |
| TdT_Index_2 | CAAGCAGAAGACGGCATACGAGATACATCGGTGACTGGAGTTCAGACGTGTGC TCTTCCGATCT | |
| TdT_Index_3 | CAAGCAGAAGACGGCATACGAGATGCCTAAGTGACTGGAGTTCAGACGTGTGC TCTTCCGATCT | |
| TdT_Index_4 | CAAGCAGAAGACGGCATACGAGATTGGTCAGTGACTGGAGTTCAGACGTGTGC TCTTCCGATCT | |
| MarSeq2 | GTCAGACCGGGGACTTATCAGCCAAC | sequencing |

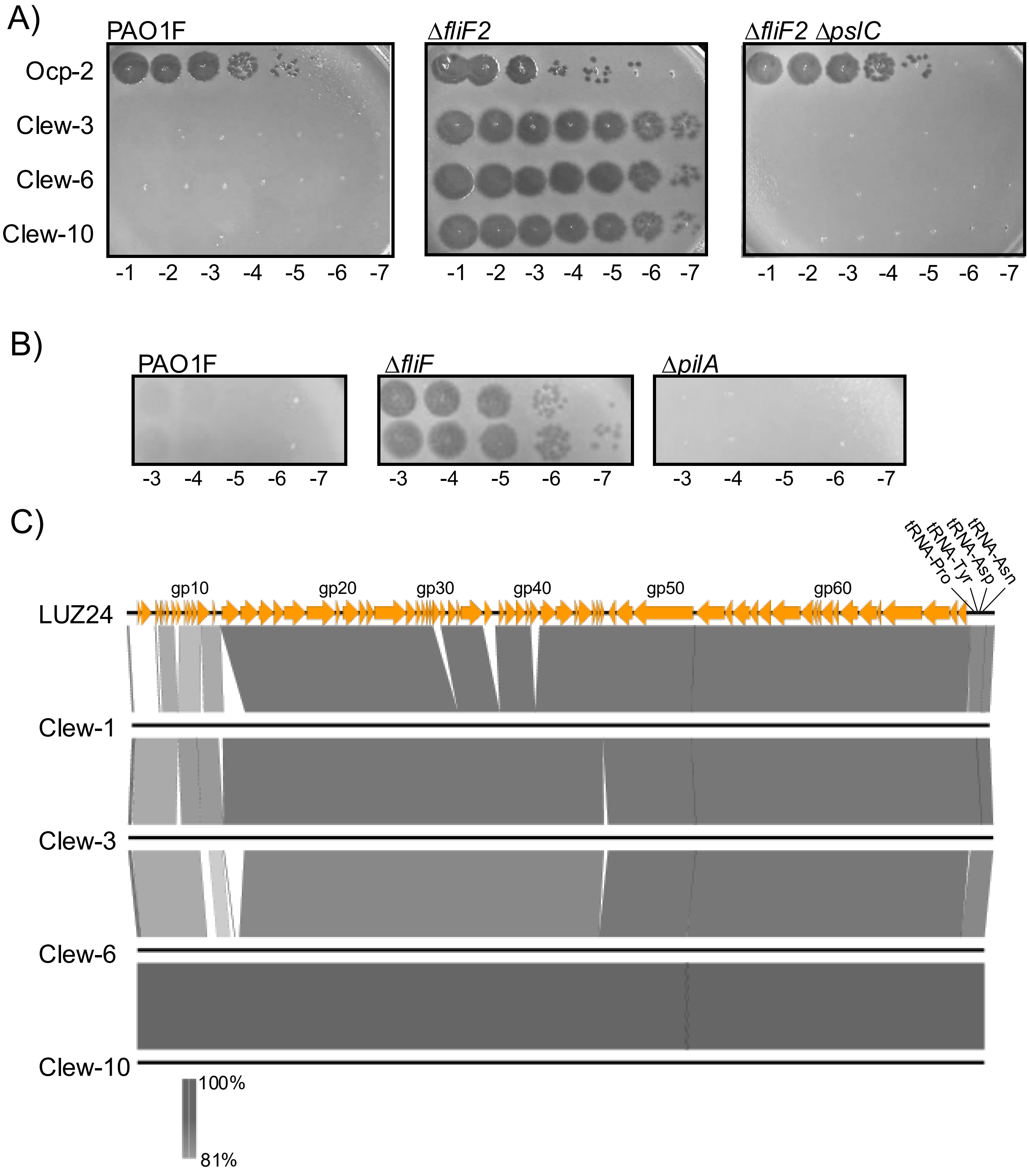


**Fig. S1 Phage Clew-3, Clew-6, and Clew-10 plaque on a *fliF* mutant of *P. aeruginosa*.** A) A dilution series of the indicated phage lysate were plated on wild type *P. aeruginosa* PAO1F, a Δ*fliF2* or Δ*fliF2* Δ*pslC* mutant strain. The data are representative of at least 2 biological replicates. B) deletion of *fliF*, not *pilA* permits phage Clew-1 to form plaques. C) Pairwise comparisons of genomes of Clew-1, Clew-3, Clew-6, and Clew-10 with Luz24, the Bruynoghevirus type strain using EasyFig. Luz24 genes are numbered sequentially and the location of every 10th ORF is indicated. (Figure generated with EasyFig)

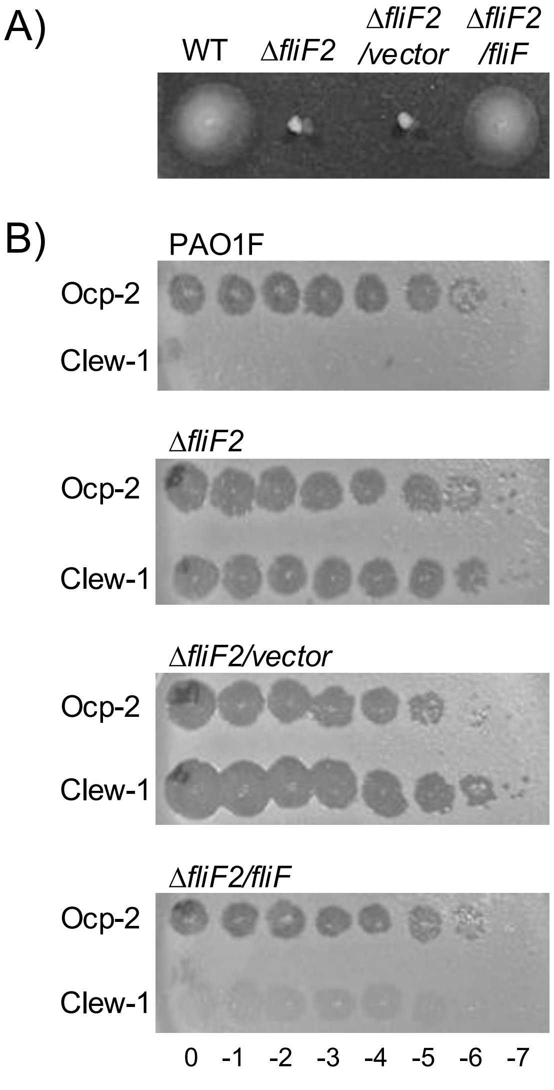


**Fig. S2. Complementation of the Δ*fliF2* mutant.** A) Swimming motility of wild type, ΔfliF2, ΔfliF2/pPSV37, or ΔfliF2/pP37-fliF assessed in 0.3% agar plates. B) Efficiency of plating experiment with the indicated strain, phage and phage dilution (log10). Data are representative of at least 3 biological replicates.

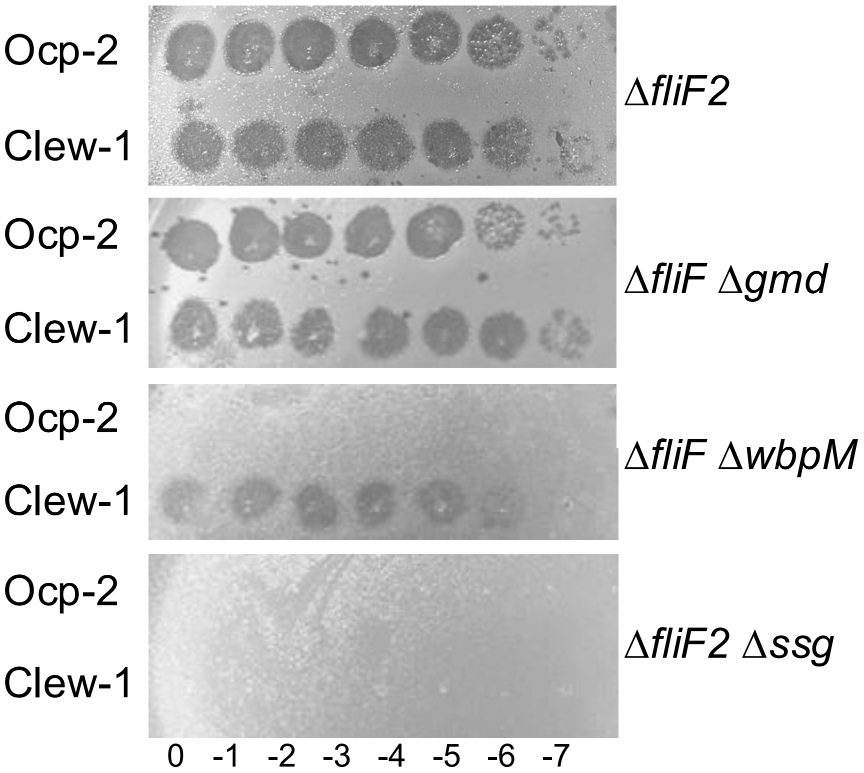


**Fig. S3. Efficiency of plating analysis on LPS mutant strains.** Ocp-2 or Clew-1 dilution series were plated on the indicated *P. aeruginosa* PAO1 mutant strains. Δ*gmd* prevents formation of A-band LPS (12), Δ*wbpM* prevents formation of B-band LPS (13), while Δ*ssg* results in a defect in the biosynthesis of the outer core (14).

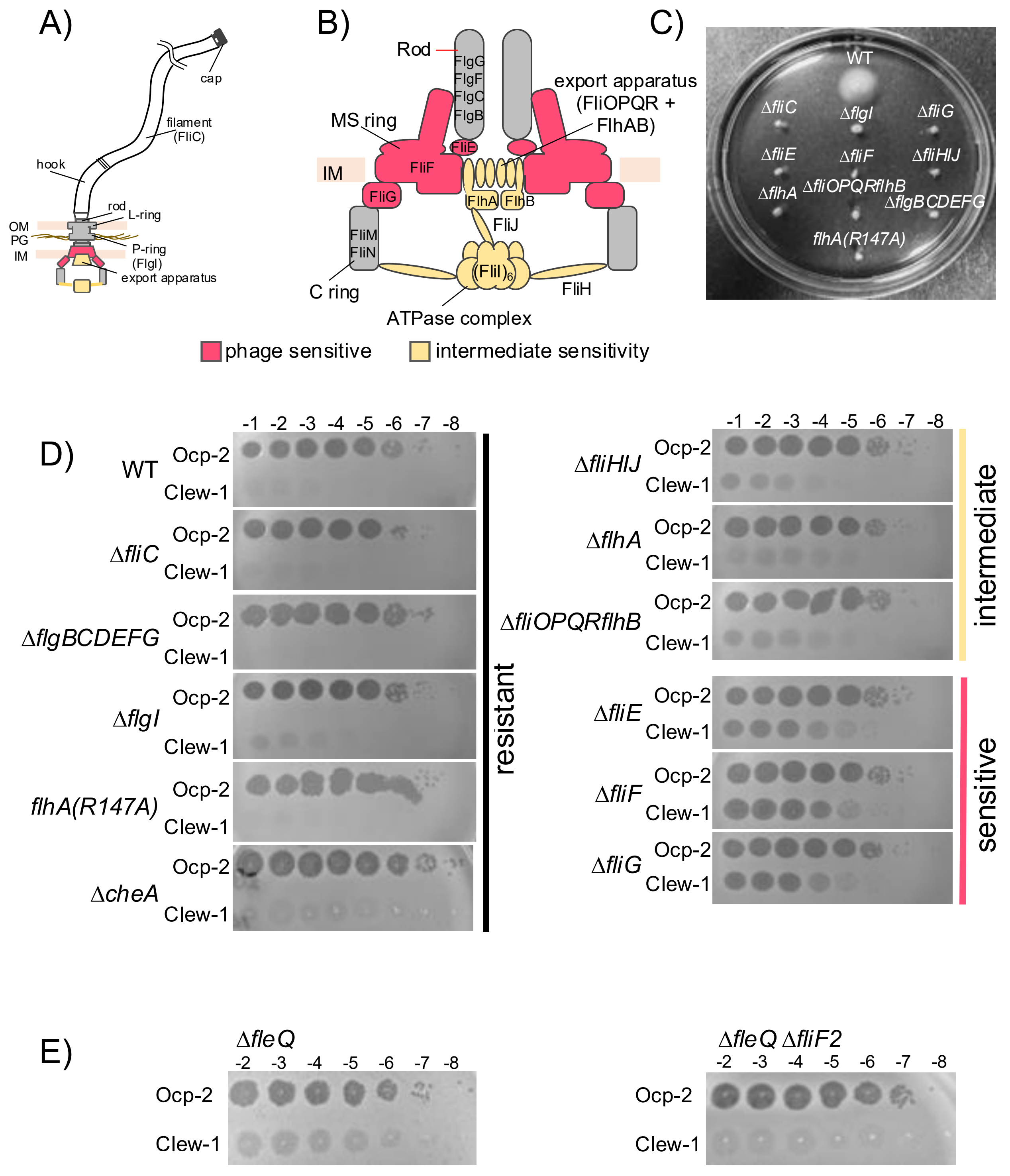


**Fig. S4. Impact of flagellar mutations on bacteriophage Clew-1 sensitivity.** Schematic of A) the flagellum and B) the flagellar basal body. Components whose removal results in complete (red) or partial (yellow) sensitivity are indicated. C) Swimming motility of wild type, as well as indicated mutant strains. D) Efficiency of plating experiment with the indicated strains, phage and phage dilutions. Data is representative of at least three biological replicates. E) Efficiency of plating experiment on *fleQ* mutant strains (representative of at least 9 biological replicates).

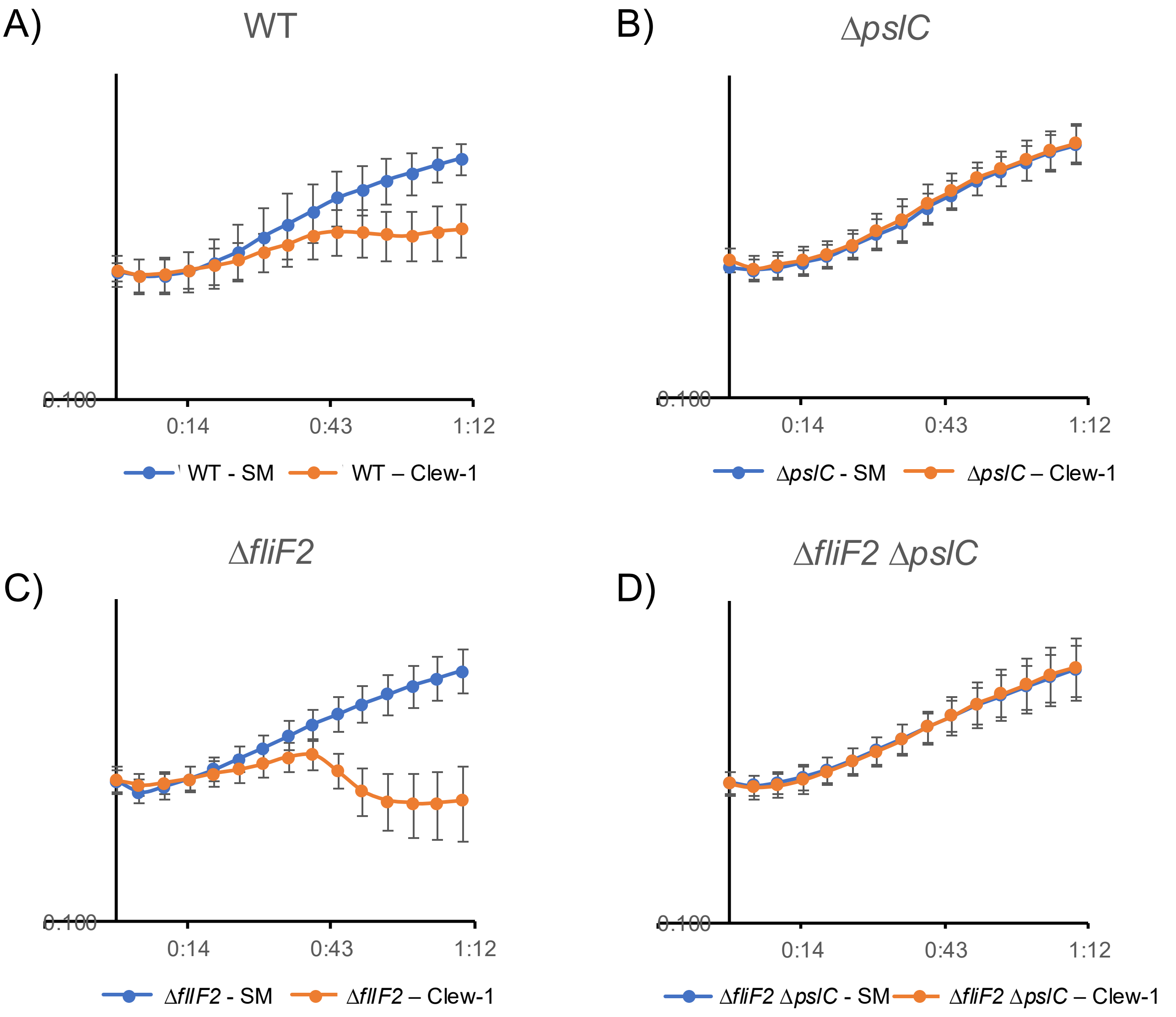


**Fig. S5. Lysis curves following bacteriophage Clew-1 infection.** Growth of A) wild type, B) Δ*pslC* mutant, C) Δ*fliF2* mutant, and D) Δ*fliF2* Δ*pslC* mutant bacteria were followed over time by monitoring the OD600 in a 96-well plate reader. Phage Clew-1 or SM-buffer were added at 0 minutes. Averages of 6 biological replicates are shown with standard deviations.

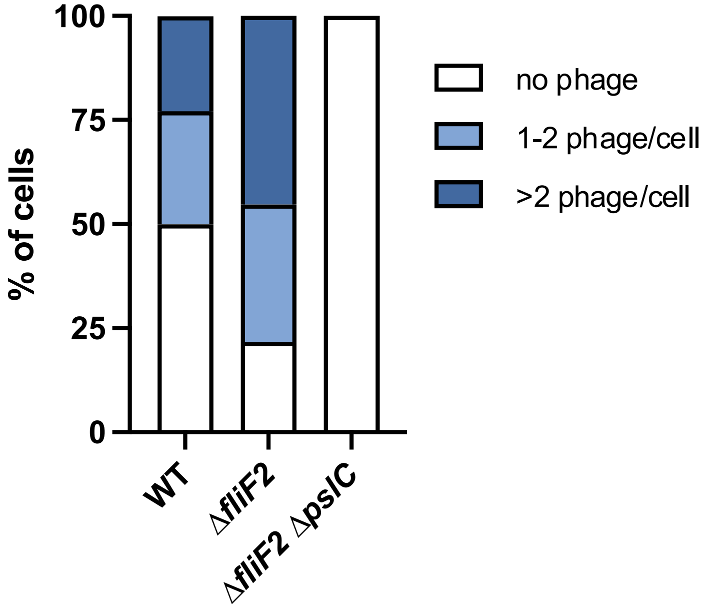


**Fig. S6. Distribution of phage attached to bacterial cells.** The phage attachment experiments in Fig. 3 was further subdivided to distinguish between bacteria harboring no attached Clew-1 phage, 1-2 phage, or more than two phage per cell. The data were analyzed by 2-way ANOVA with Šídák's multiple comparisons test. In comparing wild type and the Δ*fliF2* mutant, both the number of cells without an attached phage and the number of cells with more than 2 phage attached were significantly different ($p<0.001$ and $p<0.0001$, respectively). The difference between number of cells with 1-2 phage attached did not reach statistical significance.

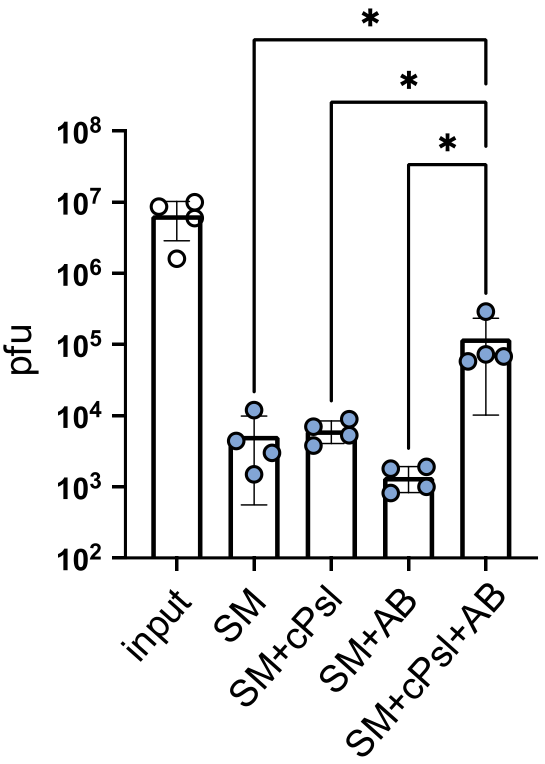


**Fig. S7. Phage Clew-1 binds to partially purified, cell-associated Psl**. Where indicated Phage Clew-1 was incubated for 1h in SM buffer with partially purified, cell-associated Psl (cPsl) and/or an anti-Psl antibody (AB). After 1h, magnetic protein-A beads were added and incubated for another 30 minutes on ice with occasional vortexing. Beads were collected and washed 3x, and phage in the input and output samples was quantified by qPCR (3 independent replicates. Statistical significance was determined by ANOVA with Sidák post-hoc test (* $p<0.05$).

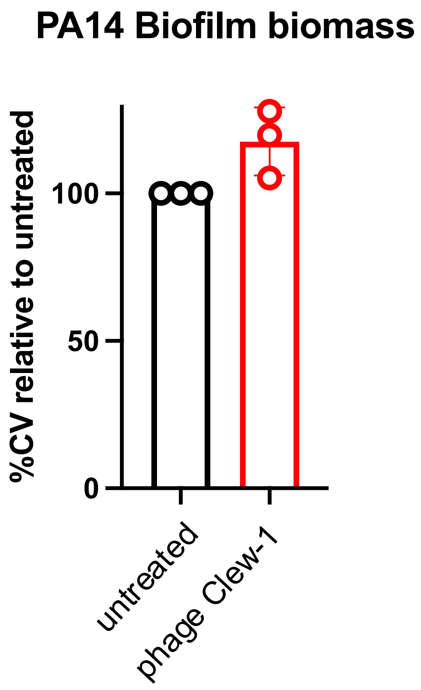


**Fig. S8.** ***P. aeruginosa*** **PA14 biofilms are insensitive to phage Clew-1.** *P. aeruginosa* PA14 biofilms were established overnight in a 96-well plate (150µL of culture, 6 technical replicates/condition), washed and incubated overnight with 200µL of LB or LB with 10^9 pfu bacteriophage Clew-1. the biofilms were then washed 3x with PBS, fixed with EtOH, dried and stained with a 0.1% crystal violet solution. The stained biofilms were washed 3x with water, and the remaining crystal violet was solubilized with 30% acetic acid. Crystal violet levels were measured spectrophotometrically at 590 nm and the untreated controls were set to 100%. Data from 3 biological replicates is shown.

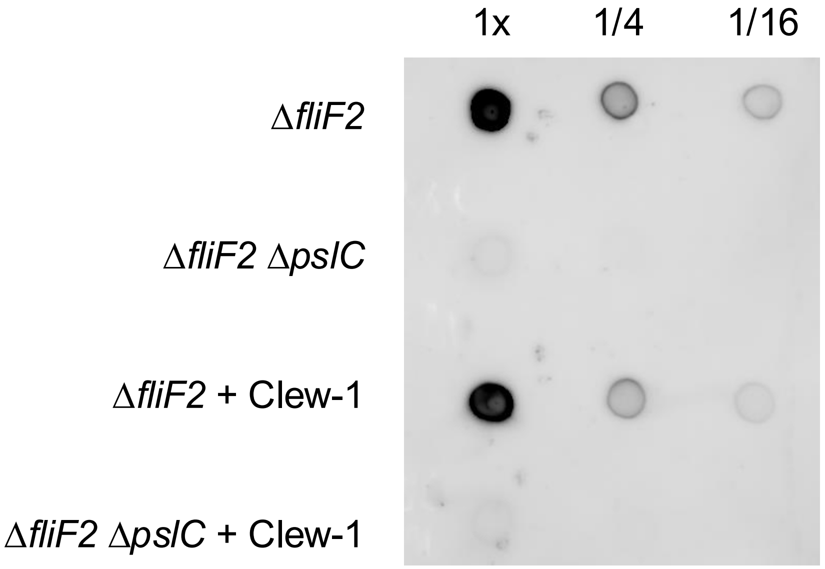


**Fig. S9. Phage Clew-1 does not to degrade Psl.** Cell free culture supernatants of PAO1F Δ*fliF2* or PAO1F Δ*fliF2* Δ*pslC* were incubated with 10^7 pfu of phage Clew-1 for 1h at 37°C (conditions that allow for phage binding and infection of *P. aeruginosa*). After the incubation, 2µL spots of the Psl+/Δpsl supernatants were spotted onto a nitrocellulose membrane, along with a 1/4 and 1/16 dilution, dried, and Psl was detected using an anti-Psl antiserum. The data are representative of 3 biological replicates.
